# Methadone Triggers Fetal Membrane Sterile Inflammation Through Innate Immune and Opioid Receptor Signaling

**DOI:** 10.64898/2026.09.25.754438

**Authors:** Megan E Kelleher, Hanah M Georges, Lily C Salmeron, Ryan W Logan, Kimberly A Yonkers, Vikki M Abrahams

## Abstract

**Introduction:** Preterm birth is often associated with fetal membrane (FM) inflammation. Opioid-use disorder during pregnancy is a major public health issue as all opioids carry a risk for preterm birth. While maintenance therapy with methadone improves maternal outcomes, it also elevates the risk of preterm birth. We previously reported that the opioid, buprenorphine, induced human FM sterile inflammation. The goal of this study was to investigate if methadone also induced FM inflammation and to determine the mechanisms involved.

**Methods:** Human FM explants were treated with or without methadone in the presence or absence of inhibitors to TLR4/MD2, TLR7, TLR8, NLRP3, the μ-opioid receptor, p65 NFκB, p38 MAPK, ERK or JNK. Supernatants were measured for cytokines/chemokines by ELISA. Tissues were analyzed by Western blot for NFκB and MAPK activity, and by RT-qPCR for the TLR7/8-activating miRs: miR-21a, miR-29a, miR-146a-3p, Let-7b. Pregnant *C57BL/6* mice were injected with PBS or methadone on E15.5 and 6hrs later FMs were analyzed by RT-qPCR.

**Results:** Methadone elevated human FM explant secretion of IL-1β, TNFα, MIP-1α, GM-CSF, and PGE2, and expression of miR-21a*. In vivo*, methadone elevated mouse FM expression of *MIP1A, KC,* and *COX2*. TLR4/MD2 inhibition had no effect on human FM inflammation in response to methadone. NLRP3 inhibition reduced methadone-induced FM IL-1β, TLR7 inhibition reduced methadone-induced FM TNFα, and inhibition of either TLR8 or the μ-opioid receptor reduced methadone-induced secretion of TNFα, MIP-1α, GM-CSF, and PGE2, but not IL-1β. Methadone increased human FM activity of p65 NFκB, p38 MAPK, JNK, and ERK and inhibition of these signaling pathways collectively reduced methadone-triggered FM inflammation. Finally, naloxone reduced most of the methadone-induced FM inflammatory response.

**Conclusion:** Methadone induces sterile inflammation in human and mouse FMs. In human FMs this inflammation is mediated through activation of innate immune TLR7, TLR8, and NLRP3; the μ-opioid receptor; and NFκB/MAPK signaling. These findings provide further insight into the mechanisms underlying the association between opioid use during pregnancy and preterm birth.

## INTRODUCTION

Opioid-use disorder (OUD) during pregnancy has increased in the US to critical levels and is a major cause of maternal morbidity and mortality (1–3). All opioids (opiate agonists) carry a risk for pregnancy complications that include preterm birth, fetal growth restriction, and fetal demise (2). The rate of preterm birth is 10.4% in the general population (4), and this more than doubles among pregnant women with untreated OUD (5). OUD also has profound neonatal consequences including neonatal opioid withdrawal syndrome (NOWS) and neurodevelopmental issues (6). Standard treatment for OUD is maintenance therapy with buprenorphine or methadone. In addition to providing a steady concentration of opioid to avoid the risk of relapse and unregulated drug use (7), treatment during pregnancy is often continued post-partum (1, 7). Despite the many benefits of treatment with buprenorphine and methadone, they are themselves associated with an elevated risk for preterm birth and NOWS (2) when compared to the general population, even if the risk might be reduced when compared to untreated OUD (8, 9). However, the mechanisms underlying OUD-associated preterm birth remain unclear.

Inflammation is a major driver of preterm birth, with infection-associated inflammation being a significant contributor (10) through activation of innate immune receptors at the maternal-fetal interface, like Toll-like receptors (TLR) and the inflammasome (11–15). However, non-infectious triggers of sterile inflammation, such as damage-associated molecular patterns (DAMPs) or dangers signals/alarmins, can also lead to prematurity by activating similar innate immune sensors in the placenta and in the fetal membranes (FM) (16). Furthermore, certain medications that are associated with preterm birth can trigger human FM sterile inflammation. We demonstrated that the serotonin reuptake inhibitor (SRI), fluoxetine, induces human FM inflammatory IL-6 secretion through p38 MAPK activation (17). We also recently reported that the opioid, buprenorphine, induces FM sterile inflammation and mediators of membrane weakening through activation of TLR4/MD2, the NLRP3 inflammasome, and the μ-opioid receptor (18). Therefore, the goal of this study was to investigate if methadone induces FM sterile inflammation and to investigate the role of innate immune and opioid receptor signaling. Herein, we report that methadone induces human FM inflammation through activation of the μ-opioid receptor, the NLRP3 inflammasome, as well as the Toll-like receptors, TLR7 and TLR8. The opioid receptor antagonist, naloxone, reduced most of this FM inflammation. This study provides further evidence for a mechanistic link between opioid use in pregnancy, FM sterile inflammation, and the elevated risk for preterm birth. Moreover, these findings may help clinicians optimize opiate agonist and antagonist treatments in pregnant women with OUD.

## MATERIALS & METHODS

### Patient Samples

Human FM tissue collection was approved by the Yale University’s Human Research Protection Program (IRB# 0607001625) and was facilitated by the Yale University Reproductive Sciences (YURS) Biobank (IRB# 1309012696) with patient consent. Human FMs were collected from uncomplicated term pregnancies (38-41 weeks’ gestation), delivered by scheduled caesarean section, in the absence of labor, known infection, or known medication/opioid use, and were transferred to the laboratory immediately after delivery.

### Animal Studies

Animal studies were approved and conducted under Yale University’s institutional animal care and use committee (Protocol #2025-11589). 8-12 week old female *C57BL/6* wildtype mice (The Jackson Lab, Bar Harbor, ME) were mated. Timed pregnant mice were injected intraperitoneally (ip) on day E15.5 with 100μl of PBS or 100μl of methadone at 10μg/kg; a dose based on prior studies (6). Mice were sacrificed 6hrs post injection. For each dam, FM tissues were dissected from all placental units, pooled, and snap frozen.

### Fetal Membrane Explants

An established FM explant system was used (18, 19). After washing FMs with PBS containing 1% penicillin (10,000 U/ml) and streptomycin (10,000 μg/ml) (Life Technologies, Grand Island, NY), and biopsied as intact 6mm explants with both the chorion and amnion layers. Areas of FMs with obvious decidual tissue attached were avoided when collecting explant biopsies. Corning Falcon 0.4μM cell culture inserts (Millipore Sigma, Burlington, MA), were placed in 24 well plates containing 1ml (500μl in the upper and lower chambers) of Dulbecco modified Eagle medium (DMEM; Life Technologies) supplemented with 10% FBS (GeminiBio), 10mM MEM non-essential amino acids, 1mM sodium pyruvate, 10mM HEPES, 100nM penicillin/streptomycin (Gibco) for overnight incubation at 37°C with 5% CO_2_. The following day, DMEM media was removed and replaced with serum-free OptiMeM media (Life Technologies). FMs were treated with either a no treatment (NT) media control or methadone (Spectrum Chemical MGF Corp, New Brunswick, NJ) at 10-200μg/ml. After establishing a dose response (Supplemental Figure 1), all subsequent *in vitro* experiments were performed with methadone at 200μg/ml. For some experiments, FMs were treated with NT or methadone in the presence of media or inhibitors to: TLR4/MD2 (LPS-RS at 10μg/ml; Invivogen, San Diego, CA); TLR7 (IRS661 at 5μM; Keck Core, Yale University) (20, 21); TLR8 (CU-CPT9a at 10μM; Invivogen); NLRP3 (MCC950 at 10μM; Invivogen); AIM2 (A151 at 5μM; Invivogen); p65 NFκB (BAY11-7085 at 10μM; Sigma Alrich, St louis, MO); p38 MAPK (SB203580 at 1μM; Selleck Chemicals, Houston, TX); ERK (SCH77298 at 10nM; Selleck Chemicals); JNK (SP600125 and 25μM; Sigma Aldrich); or the μ-opioid receptor (β-FNA at 10μM; Sigma Aldrich). FMs were also treated with NT or methadone in the presence of media or (-) naloxone (80μg/ml; Thomas Scientific, Swedesboro, NJ). Inhibitors or (-) naloxone were added to FM culture system 1hr prior to methadone treatment. After 48hrs, cell-free supernatants and FM tissues were collected. Tissues were snap frozen. Supernatants were centrifuged to remove any cellular debris. Both tissues and supernatants were stored at -80°C. All FM experiments were performed in triplicate.

### Human Fetal Membrane Secreted Factors

Supernatants collected from human FM explants were measured by ELISA for IL-1β, IL-6, IL-8, TNFα, G-CSF, GM-CSF, MIP-1α, PGE2, MMP1, MMP2, MMP9 and TIMP2 (R&D Systems, Minneapolis, MN).

### Western Blot Analysis

Prior to western blot analysis, FM tissues were homogenized for protein in lysis buffer containing phosphatase inhibitors (Cell Signaling Technology, Danvers, MA) and protein concentrations were measured using the Pierce BCA protein assay (ThermoFisher Scientific, Waltham, MA). Following SDS-PAGE gel electrophoresis and subsequent transfer to PVDF membranes (Bio-Rad, Hercules, CA), proteins were probed for using following primary antibodies all from Cell Signaling Technology: phosphorylated (p)-p65 NFκB (#3033); total (t)-p65 NFκB (#8242); p-p38 MAPK (#9211); t-p38 MAPK (#9212); p-ERK (#9101); t-ERK (#4695); p-JNK (#9251); t-JNK (#9252). Western blot images were captured using an Amersham Imager 680 (General Electric, Boston, MA). Image Studio Lite (Li-Cor Biosciences, Lincoln, NE) was used to perform semi-quantitative densitometry.

### RT-qPCR

Human and mouse FM tissues were homogenized using a beadbug homogenizer with microtubes pre-filled with high impact zirconium beads (Benchmark Scientific; Sayreville, NJ) as previously described (20). RNA was extracted using TRIzol (ThermoFisher Scientific) according to manufacturer’s protocol. RNA concentrations were measured using a NanoDrop 2000 Microvolume Spectrophotometer (ThermoFisher Scientific). Human FM tissue expression of miR-21a, miR-29a, miR-146a-3p, and Let-7b were measured using the Taqman MicroRNA Assay (Life Technologies) according to manufacturer’s protocols for reverse transcription and qPCR. For miR-146a-3p, a pre-amplification step was included prior to performing PCR using the Taqman PreAmp master mix kit (Life Technologies). Small nuclear RNA U6 was used as a reference gene for the normalization of miR expression. Mouse FM expression of *IL1B, TNFA, KC, MIP1A and COX2* was measured by RT-qPCR using the SuperScript II kit (Invitrogen; Waltham, MA) and the Kapa SYBR Fast qPCR kit (Sigma-Aldrich) and normalized to *ACTIN*. RT-qPCR data was analyzed using the 2^−ΔΔCT^ method and reported as either fold change (FC) or relative abundance (RA).

### Bulk RNA-Seq Analysis

#### RNA Seq Quality Control

RNA samples from human FM tissues were submitted to the Yale Center for Genome Analysis (YCGA) for RNA sequencing. Total RNA quality was determined by estimating the A260/A280 and A260/A230 ratios by nanodrop. RNA integrity was determined by running an Agilent Bioanalyzer gel, which measures the ratio of the ribosomal peaks.

#### RNA Seq Library Prep

Using the Kapa RNA HyperPrep Kit with RiboErase (KR1351), rRNA was depleted starting from 25-1000ng of total RNA by hybridization of rRNA to complementary DNA oligonucleotides, followed by treatment with RNase H and DNase to remove rRNA duplexed to DNA. Samples were then fragmented using heat and magnesium. 1^st^ strand synthesis was performed using random priming. 2^nd^ strand synthesis incorporated dUTPs into the 2^nd^ strand cDNA. Adapters were then ligated and the library was amplified. Strands marked with dUTPs were not amplified allowing for strand-specific sequencing. Indexed libraries that met appropriate cut-offs for both quantity and quality were quantified by qRT-PCR using a commercially available kit (KAPA Biosystems) and insert size distribution was determined with the LabChip GX or Agilent Bioanalyzer. Samples with a yield of ≥0.5 ng/ul were used for sequencing.

#### Flow Cell Preparation and Sequencing

Sample concentrations were normalized to 2 nM and loaded onto an Illumina NovaSeq X flow cell at a concentration that yields 25 million passing filter clusters per sample. Samples were sequenced using 100bp paired-end sequencing on an Illumina NovaSeq X according to Illumina protocols. The 10bp unique dual index was read during additional sequencing reads that automatically follow the completion of read 1. Data generated during sequencing runs were simultaneously transferred to the YCGA high-performance computing cluster. A positive control (prepared bacteriophage Phi X library) provided by Illumina is spiked into every lane at a concentration of 0.3% to monitor sequencing quality in real time.

#### Data Analysis and Storage

Signal intensities were converted to individual base calls during a run using the system’s Real Time Analysis (RTA) software. Base calls were transferred from the machine’s dedicated personal computer to the Yale High-Performance Computing cluster via a 10 Gigabit network mount for downstream analysis. Primary analysis (sample de-multiplexing) was performed using bcl2fastq2. The data were used if the sample error rate was less than 2% and the distribution of reads per sample in a lane was within reasonable tolerance. Aligned reads were analyzed for differential gene expression in R using the Bioconductor package DESeq2 with multiple comparison adjustments made using the Benjamini-Hochberg method. Following differential expression analysis, gene pathways and interactions were analyzed using Ingenuity Pathways Analysis (IPA; Qiagen Hilden, Germany). Raw and processed data are archived in the NCBI GEO database: Accession #: GSE346101.

### Statistical analysis

For each human FM tissue sample, explant treatments were performed in three technical replicates, pooled, and reported as one experiment. Experiments were performed in four or more biological/independent replicates, with sample size reported in the figure legends as “n=”. Analysis was performed in duplicate and averaged, and data are reported as mean ± SEM of independent experiments with statistical significance defined as *p*<0.05. Statistical analysis was performed using GraphPad Prism (GraphPad Software, La Jolla, CA). For normally distributed data, data were analyzed using either one-way analysis of variance (ANOVA) for multiple comparisons or a paired t-test. For data not normally distributed, data were analyzed using a non-parametric multiple comparison test or the Wilcoxon matched-pairs signed rank test. Outliers were identified using the ROUT method Q coefficient of 1%.

## RESULTS

### Methadone triggers FM sterile inflammation

The impact of methadone on FMs was first assessed by measuring the effect of various doses of methadone (0-200μg/ml) on human FM explant secretion of a range of inflammatory factors (IL-1β, IL-6, IL-8, TNF-α, G-CSF, GM-CSF, MIP-1α), mediators of uterine contractions (PGE2), and mediators of membrane weakening (MMP1, MMP2, MMP9 These factors have been associated with preterm birth (22–25), and most were previously shown to be elevated in human FM explants treated with buprenorphine (18). As shown in Figure 1A and Supplemental Figure 1, methadone at 200μg/ml optimally elevated human FM explant secretion of IL-1β, TNF-α, GM-CSF, MIP-1α and PGE2 when compared to the no treatment (NT) or 0μg/ml controls. Methadone had no effect on FM secretion of inflammatory IL-6, IL-8, or G-CSF (Supplemental Figure 1). Human FM explant secretion of the mediators of membrane weakening, MMP1 and MMP9, were also unchanged by methadone exposure (Supplemental Figure 1). Treatment of human FM explants with methadone significantly reduced MMP2 secretion (Supplemental Figure 1), while having no effect on its inhibitor, TIMP2 (Supplemental Figure 2).

**Figure 1.**
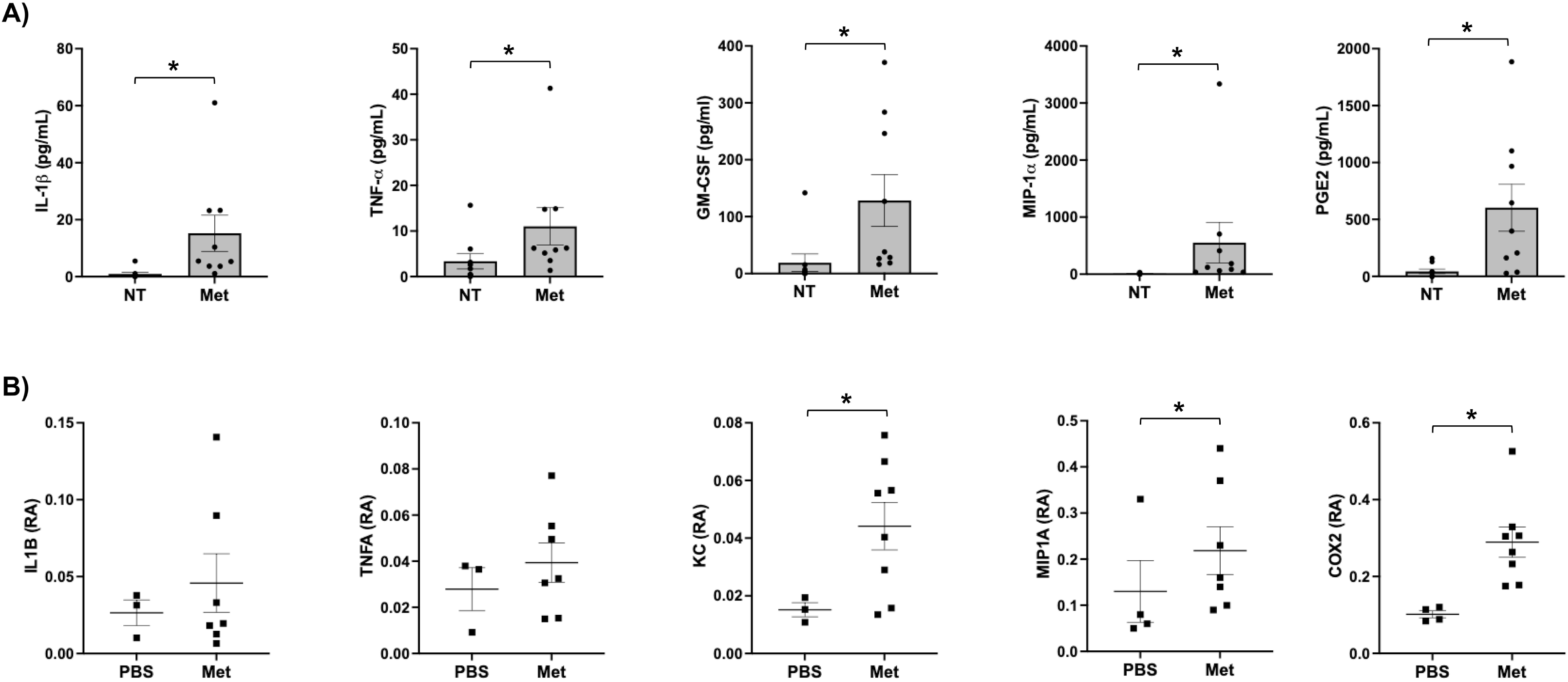
Methadone triggers FM sterile inflammation. (A) Human FM explants were treated with no treatment (NT) or Methadone (Met) at 200μg/ml (n=6-9). After 48hrs, cell-free supernatants were collected and measured by ELISA for IL-1β, TNFα, GM-CSF, MIP-1α, and PGE2 (\**p*<0.05). (B) 8-12 week old female C57BL/6 wildtype timed pregnant mice were injected intraperitoneally (ip) on day E15.5 with PBS (n=4) or methadone (Met) (n=8) and were sacrificed 6hrs later. FM tissues were dissected from the placental unit, homogenized for RNA, and RT-qPCR used to measure *IL1B*, *TNFA*, *KC*, *MIP1A*, and *COX2*. Data are presented as relative abundance (RA) of expression (\**p*<0.05).

In support of our *in vitro* findings using human FM explants, which models acute opioid exposure (Figure 1A and Supplemental Figure 2), FMs collected from pregnant mice with an acute exposure to methadone at E15.5, 6hrs later showed a significant increase in *MIP1A* when compared to the PBS controls (Figure 1B). Similarly, methadone significantly increased FM expression of *COX2* (Figure 1B) which is used as a marker of PGE2 production. Methadone exposure also significantly increased mouse FM inflammatory *KC*, and there was a trend for elevated *IL1B* and *TNFA* in response to methadone (Figure 1B), again aligning with our *in vitro* data (Figure 1A).

When RNA-seq was performed on human FM tissues that were either untreated or exposed to methadone, one of the top significant pathways highlighted from IPA was “*Cytokine Storm*” (Figure 2A and Supplemental Figure 3). Within this pathway, and within the top 10 genes found to be significantly unregulated in response to methadone, was GM-CSF (CSF2) and TNFA (Figure 2B), validating our ELISA data (Figure 1A). IPA analysis also revealed increased inflammasome-associated IL1 (Figure 2A and Supplemental Figure 3). Additionally, IPA revealed a number of genes in this pathway to be downregulated (Figure 2A and Supplemental Figure 3).

**Figure 2.**
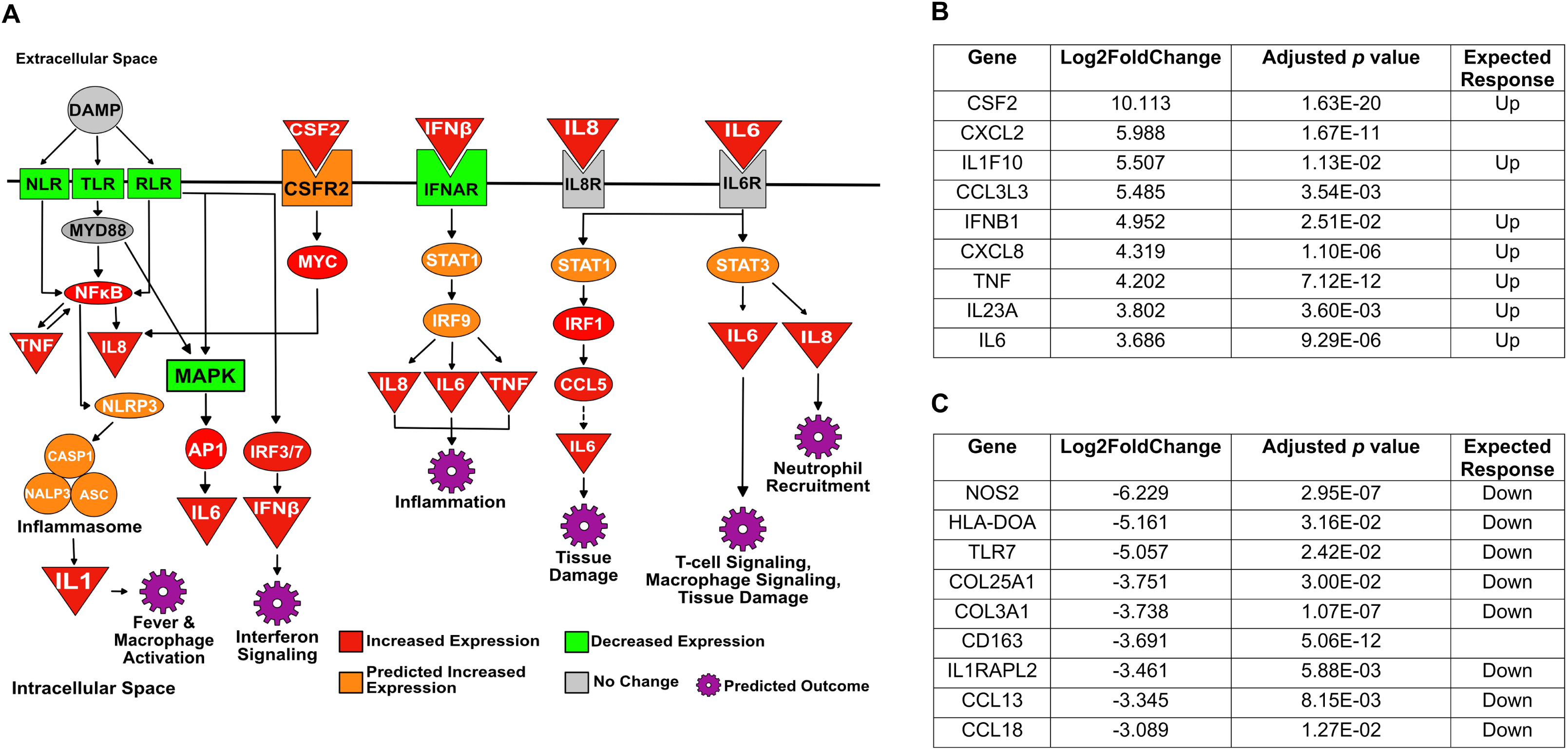
RNA-seq analysis. Human FM explants were treated with no treatment (NT) or Methadone (Met) at 200μg/ml (n=6). After 48hrs total FM tissue RNA was extracted and bulk RNA-seq analysis performed. (A) IPA analysis revealed “Cytokine Storm” in the top 10 pathways. (B) Table shows the top 10 upregulated genes from the “Cytokine Storm” IPA. (C) Table shows the top 10 downregulated genes from the “Cytokine Storm” IPA.

### Methadone-induced human FM inflammation is independent of TLR4/MD2 activation

In non-pregnant systems, methadone can activate the innate immune receptor, TLR4, by interacting with its adapter protein, MD2, leading to inflammation (26). Thus, this signaling pathway was investigated in the human FM sterile inflammatory response to methadone. As shown in Figure 3A, the TLR4/MD2 antagonist, LPS-RS, was unable to reverse the effects on methadone on FM secretion of IL-1β, TNF-α, GM-CSF, MIP-1α or PGE2.

**Figure 3.**
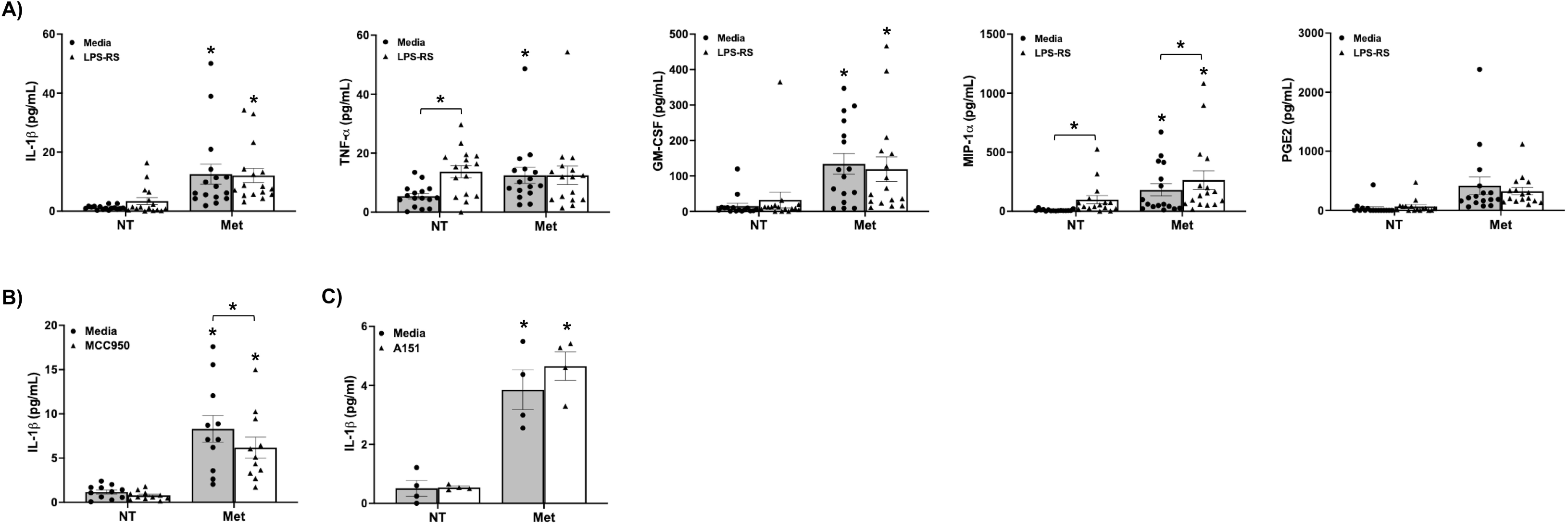
Methadone-induced human FM inflammation is independent of TLR4/MD2 activation, but the IL-1β response is dependent on the NLRP3 inflammasome. (A) Human FM explants were treated with no treatment (NT) or Methadone (Met) in the presence of media of the TLR4/MD2 inhibitor, LPS-RS (n=16). After 48hrs, cell-free supernatants were collected and measured by ELISA for IL-1β, TNFα, GM-CSF, MIP-1α, and PGE2. \**p*<0.05 relative to the NT/Media or NT/inhibitor control. (B-C) Human FM explants were treated with no treatment (NT) or Methadone (Met) in the presence of media or (B) the NLRP3 inhibitor, MCC950 (n=11) or (C) the AIM2 inhibitor, A151 (n=4). After 48hrs, cell-free supernatants were collected and measured by ELISA for IL-1β. \**p*<0.05 relative to the NT/Media or NT/inhibitor control unless otherwise indicated.

### Methadone-induced human FM IL-1β secretion is partially dependent on the NLRP3 inflammasome

Activation of the NLRP3 inflammasome plays an important role in FM production of IL-1β (19). As shown in Figure 3B, the NLRP3 inhibitor, MCC950, partially reduced methadone-induced human FM explant IL-1β secretion by 23.8±5.8%. In contrast, the AIM2 inflammasome inhibitor, A151, had no effect on the ability of methadone to trigger FM IL-1β secretion (Figure 3C).

### Methadone-induced human FM inflammation is dependent on activation of the μ-opioid receptor

While methadone can weakly agonize the κ-opioid receptor, it is a particularly strong, long acting, agonist of the μ-opioid receptor. The specific μ-opioid receptor antagonist, β-FNA, significantly reduced the ability of methadone to increase human FM secretion of all inflammatory factors (TNF-α, GM-CSF, MIP-1α, PGE2) with the exception of inflammasome-mediated IL-1β (Figure 4A). Specifically, β-FNA significantly reduced methadone triggered FM secretion of TNF-α by 49.2±11.8%, GM-CSF by 38.9±10.0%, MIP-1α by 45.6±5.9%, and PGE2 by 50.7±11.5% (Figure 4A). Similarly, (-) naloxone, which is a broad opioid receptor antagonist (27), also significantly reduced the ability of methadone to increase human FM secretion of TNF-α by 36.1±10.8%, GM-CSF by 41.5±13.7%, and MIP-1α by 37.5±10.3%, again without affecting IL-1β secretion (Figure 4B). Methadone-induced FM secretion of PGE2 also trended to being reduced by naloxone by 26.1±7.3 (*p*=0.08) (Figure 4B).

**Figure 4.**
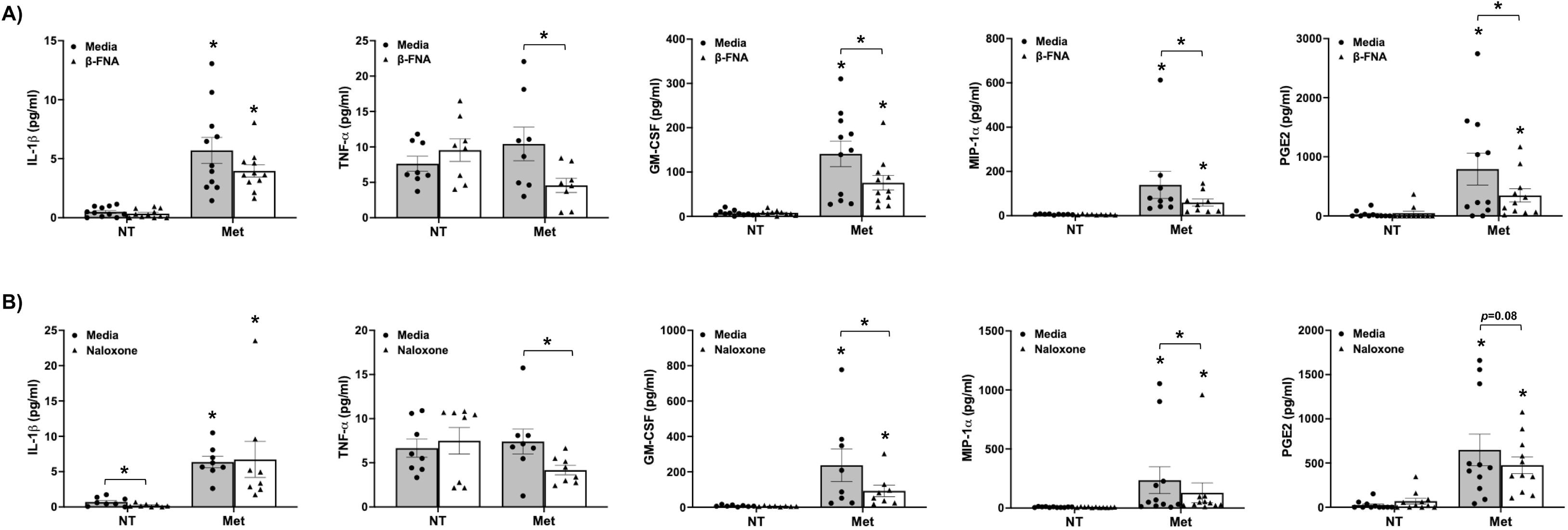
Methadone-induced human FM inflammation is dependent on activation of the μ-opioid receptor. Human FM explants were treated with no treatment (NT) or Methadone (Met) in the presence of media or (A) the μ-opioid receptor antagonist, β-FNA (n=8-11) and (B) naloxone (n=8-11). After 48hrs, cell-free supernatants were collected and measured by ELISA for IL-1β, TNFα, GM-CSF, MIP- 1α, and PGE2. \**p*<0.05 relative to the NT/Media or NT/inhibitor control unless otherwise indicated.

### Methadone-induced human FM inflammation is dependent on activation of TLR7 and TLR8

Having demonstrated that a combination of μ-opioid receptor and NLRP3 inflammasome signaling mediates methadone-induced human FM inflammation, we next asked what downstream mechanisms might be involved. We recently reported in human FMs that downstream of TLR activation by infectious components, the TLR7/TLR8-activating microRNA, miR-146a-3p (21, 28), acts as an intermediate secondary signal that subsequently activates TLR7 and TLR8, contributing to the FM inflammatory response (20, 29, 30). So far, miR-21a-5p, miR29a-5p, miR146a-3p and Let-7b have been identified and characterized as miRs that can specifically activate TLR7 and/or TLR8 (21, 28, 31). Thus, we sought to determine whether this novel signaling mechanism might be involved in opioid-induced FM sterile inflammation. As shown in Figure 5A, methadone significantly elevated human FM tissue expression of miR-21a without significantly affecting the levels of miR-29a, miR146a-3p or Let-7b. Inhibition of TLR7 signaling by the inhibitor, IRS661, significantly reduced methadone-induced FM secretion of TNF-α by 50.1±14.3% without affecting IL-1β, GM-CSF, MIP-1α or PGE2 (Figure 5B). The TLR8 inhibitor, CU-CPT9a, significantly reduced the ability of methadone to elevate FM secretion of TNF-α by 39.5±8.0%, GM-CSF by 51.8±10.1%, and PGE2 by 52.9±14.1% (Figure 5C). FM secretion of MIP-1α in response to methadone also trended to being reduced by CU-CPT9a by 41.3±9.9% (*p*=0.08).

**Figure 5.**
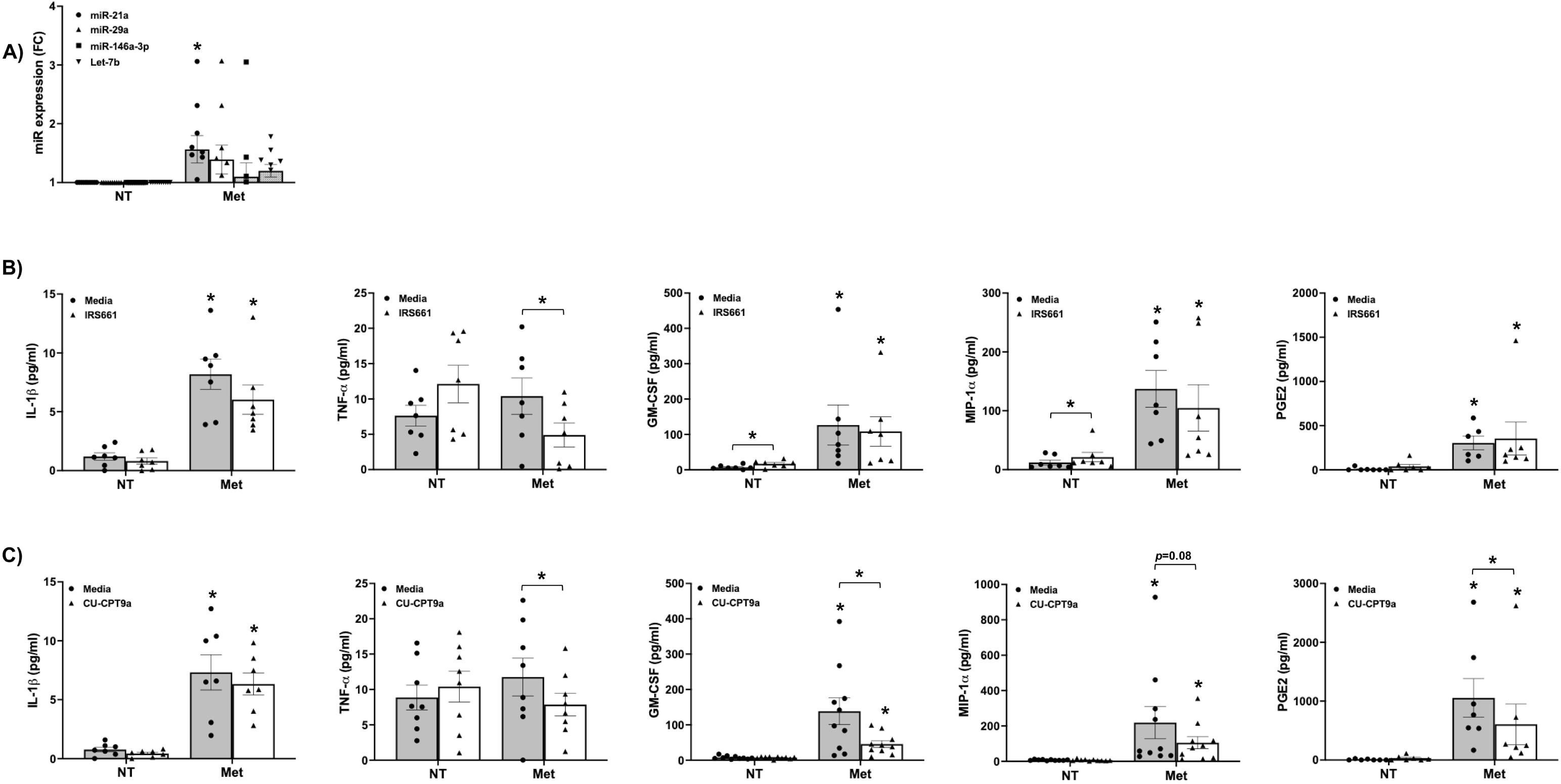
Methadone-induced human FM inflammation is dependent on activation of TLR7 and TLR8. (A) Human FM explants were treated with no treatment (NT) or Methadone (Met) at 200μg/ml (n=10). After 48hrs tissues were homogenized for RNA and RTqPCR performed to measure expression of the TLR7/TLR8-activating miRs: miR-21a, miR-29, miR-146a-3p and Let-7b. Data are expression as fold change (FC) relative to the NT control (\**p*<0.05 *vs.* the NT control). (B-C) Human FM explants were treated with no treatment (NT) or Methadone (Met) in the presence of media or (B) the TLR7 inhibitor, IRS661 (n=7) and (C) the TLR7 inhibitor, CU-CPT9a (n=7-10). After 48hrs, cell-free supernatants were collected and measured by ELISA for IL-1β, TNFα, GM-CSF, MIP-1α, and PGE2. \**p*<0.05 relative to the NT/Media or NT/inhibitor control unless otherwise indicated.

### Methadone-induced human FM inflammation is dependent on the p65 NFκB, p38 MAPK, ERK, and JNK signaling pathways

Activation of TLRs trigger downstream NFκB and MAPK pathways (32), and opioid receptors can activate downstream p38 MAPK and ERK signaling (33). As shown in Figure 6, methadone exposure significantly elevated human FM explant phosphorylation of p65 NFκB, p38 MAPK, ERK and JNK, indicating activation of these signaling pathways. To determine the contribution of these signaling pathways to methadone-induced human FM inflammation, specific inhibitors were utilized. As shown in Figure 7, FM secretion of TNF-α and MIP-1α in response to methadone was reduced by inhibitors to p65 NFκB, p38 MAPK, ERK and JNK. Methadone-triggered GM-CSF secretion was reduced by inhibitors to p65 NFκB, p38 MAPK, and ERK, but not JNK (Figure 7). FM IL-1β secretion in response to methadone was reduced by inhibitors to ERK and JNK, but not p65 NFκB or p38 MAPK, and FM PGE2 secretion in response to methadone was only reduced by the p38 MAPK inhibitor (Figure 7).

**Figure 6.**
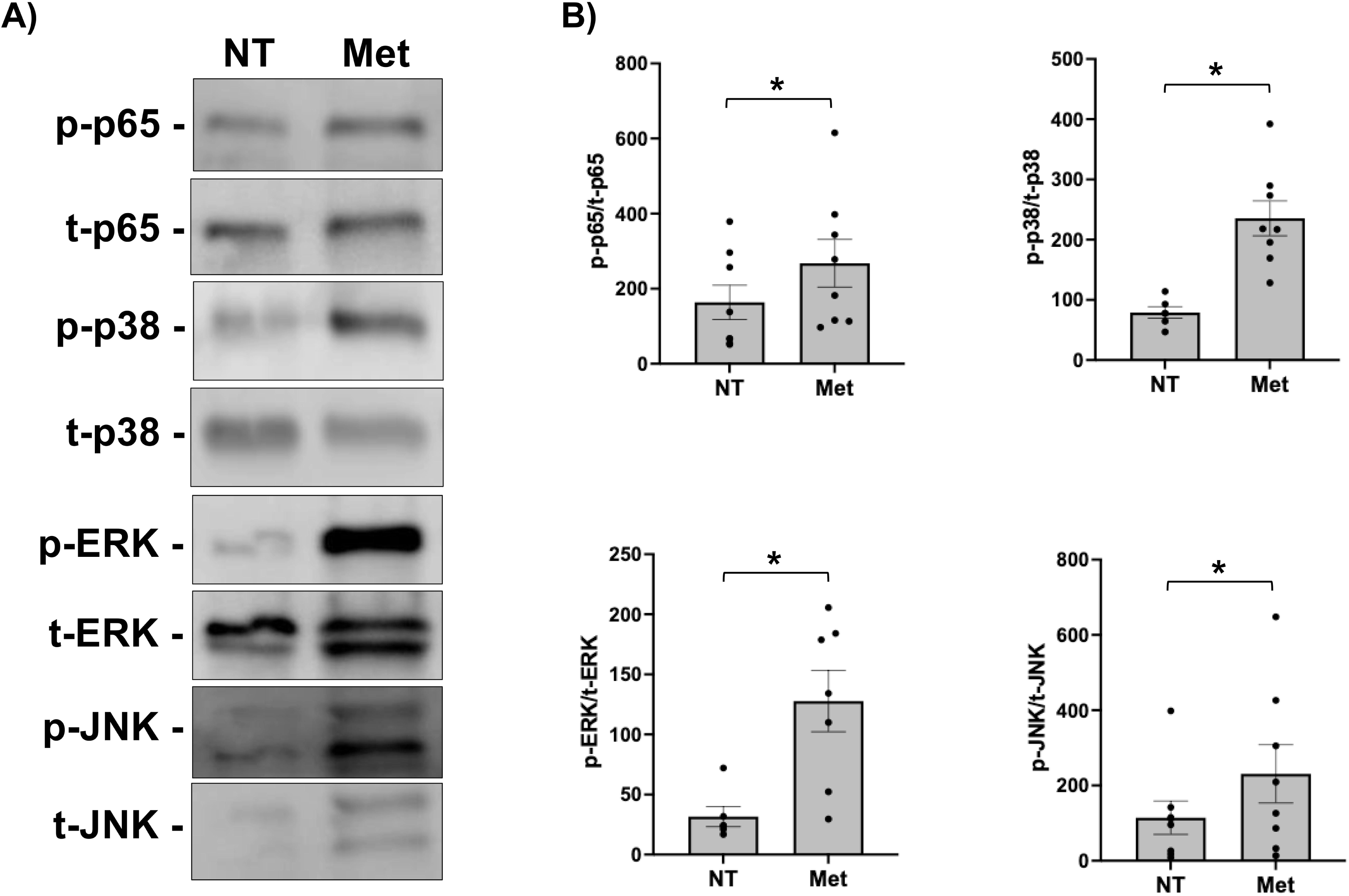
Methadone activates human FM p65 NFκB, p38 MAPK, ERK and JNK signaling pathways. Human FM explants were treated with no treatment (NT) or Methadone (Met) (n=8). After 48hrs tissues were homogenized for protein and Western Blot Analysis performed for phosphorylated (p) and total (t) . p65 NFκB, p38 MAPK, ERK and JNK. (A) Western Blot images are from representative blots. (B) Bar charts show quantification of protein expression as determined by densitometry and expressed as the ratio of signals for phosphorylated (p) to total (t) protein (\**p* < 0.05).

**Figure 7.**
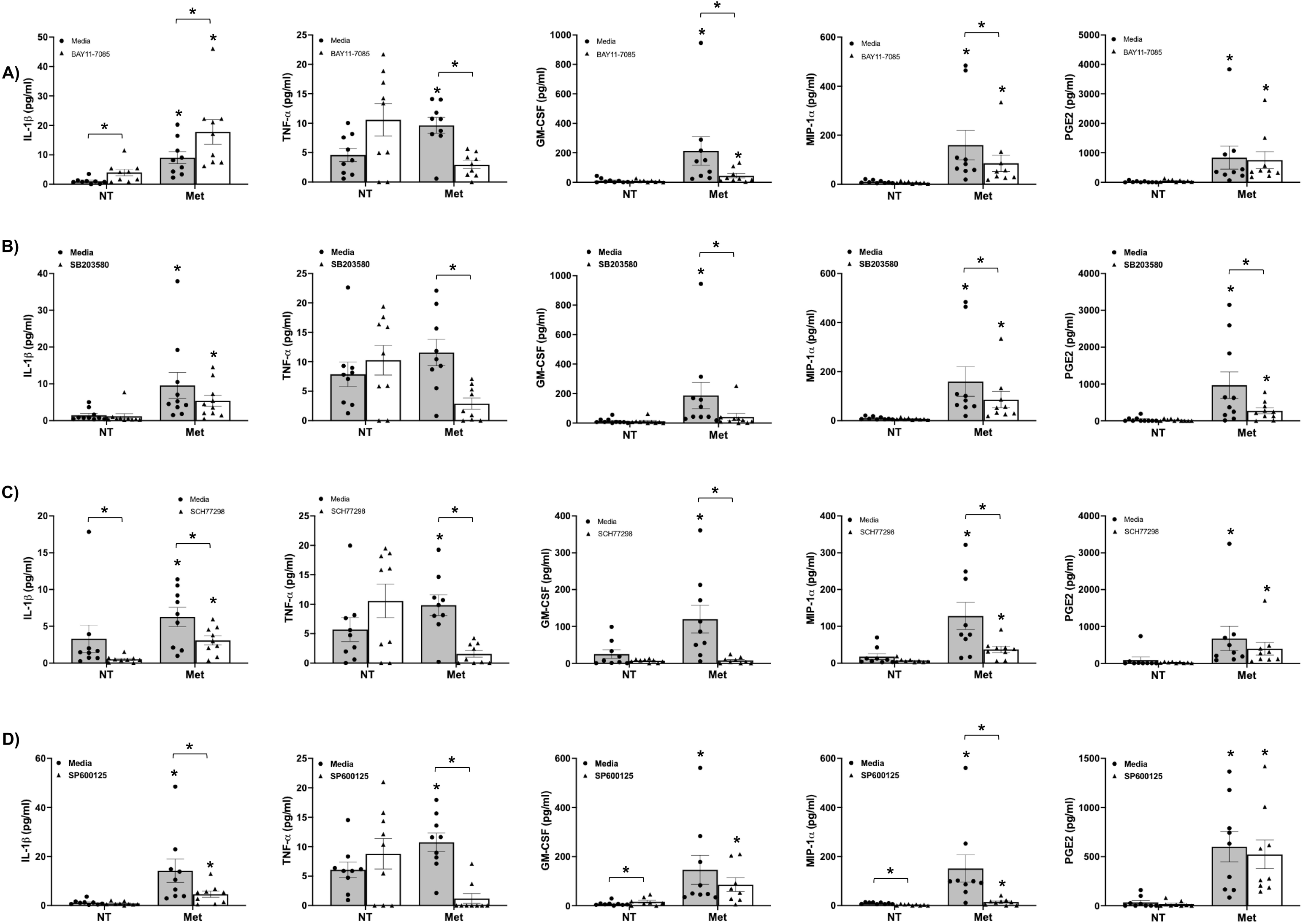
Methadone-induced human FM inflammation is dependent on the p65 NFκB, p38 MAPK, ERK, and JNK signaling pathways. Human FM explants were treated with no treatment (NT) or Methadone (Met) in the presence of media or (A) the p65 NFκB inhibitor, BAY11-7085 (n=9), (B) the p38 MAPK inhibitor, SB203580 (n=9-10), (C) the ERK inhibitor, SCH77298 (n=9), and the (D) the JNK inhibitor, SP600125 (n=9). After 48hrs, cell-free supernatants were collected and measured by ELISA for IL-1β, TNFα, GM-CSF, MIP-1α, and PGE2. \**p*<0.05 relative to the NT/Media or NT/inhibitor control unless otherwise indicated.

## DISCUSSION

Inflammation is a common underlying cause of preterm birth (10). We and others have found that infectious and non-infectious insults induce inflammatory responses in FMs that includes cytokines and chemokines associated with preterm birth, like TNFα, IL-6 and inflammasome-mediated IL-1β (17, 19, 34–38), and there is evidence of elevated inflammasome activation in FMs from women with preterm birth (39).

In non-pregnant systems, the medications for OUD (MOUD), methadone and buprenorphine, can trigger inflammation through activation of TLR4/MD2 and opioid receptors (26). Despite the association between opioid use during pregnancy and an increased risk of preterm birth (2), little is still known about how these medications impact pregnancy. In this current study, we report that methadone triggers human FM sterile inflammation through activation of innate immune TLR7, TLR8, and the NLRP3 inflammasome, as well as the μ-opioid receptor, and downstream NFκB and MAPK signaling (Table 1).

**Table 1:** Summary of human FM responses to methadone and the mechanisms involved. (✓) Involved; (⌋) Not involved; (-) Not tested.

| | TLR4/MD2 | TLR7 | TLR8 | NLRP3 | μ-opioid<br>receptor | P65 NFκB | P38 MAPK | ERK | JNK |
| --- | --- | --- | --- | --- | --- | --- | --- | --- | --- |
| IL-1β | x | x | x | ✓ | x | x | x | ✓ | ✓ |
| TNF-α | x | ✓ | ✓ | - | ✓ | ✓ | ✓ | ✓ | ✓ |
| GM-CSF | x | x | ✓ | - | ✓ | ✓ | ✓ | ✓ | x |
| MIP-1 $\alpha$ | x | x | ✓ | - | ✓ | ✓ | ✓ | ✓ | ✓ |
| PGE2 | x | x | ✓ | - | ✓ | x | ✓ | x | x |

In our initial published study using buprenorphine (18), and in this current follow-up study using methadone, we found both opioids induce human FM inflammation by elevating the secretion of IL-1β, TNFα, GM-CSF, MIP-1α, and PGE2; factors associated with preterm birth (22–25). However, in contrast to our findings with methadone, buprenorphine also increased FM secretion of additional inflammatory factors also associated with preterm birth like IL-6, IL-8 and G-CSF, as well as the mediators of membrane weakening, MMP1and MMP9 (18). In our current study methadone did not elevate FM secretion of MMP1 or MMP9 and reduced basal MMP2 levels. Based on this differential membrane weakening response, it would be of interest to determine whether buprenorphine carries a higher risk for premature rupture of membranes (PROM) than methadone. While opioids elevate the risk for PROM, clinical studies have not stratified based on specific opiate agonists (40, 41).

In addition to the human FM inflammatory profiles generated in response to buprenorphine and methadone overlapping, yet distinct, there was also a difference in the magnitude of responses generated as methadone induced a stronger FM IL-1β, MIP-1α and PGE_2_ response, when compared to buprenorphine (18). While the dose of methadone used in our *in vitro* studies appears high compared to that in the circulation, there is evidence from an *in vivo* mouse model and an *ex vivo* human placental perfusion study that methadone is retained in the placenta where it accumulates at high levels that may prolong its action and delay clearance (6, 42). Similar to our studies with buprenorphine (18), RNA-seq analysis on FM tissues exposed to methadone validated our protein-based findings that methadone, like buprenorphine, triggered a strong FM inflammatory response - a “cytokine storm”. Inflammatory factors already identified as being upregulated in FMs in response to methadone by ELISA were validated by our RNA-seq data and IPA analysis (IL1, TNF, CSF2/GM-CSF). However, transcriptional increases in IL-6 and CXCL8/IL-8, were not seen at the protein level, which could be due to the sensitivity of RNA-seq or that changes in RNA expression do not always translate into changes at the protein level.

In support of our *in vitro* findings in human FMs, which models acute opioid exposure, FMs collected from wildtype pregnant mice with an acute exposure to methadone at E15.5, 6hrs later showed a significant increase in *MIP1A*, *KC* and *COX2.* Similarly, methadone significantly increased FM expression of *COX2*, a marker of PGE2 production that is increased in mouse models of preterm birth, including in FMs (43), and in preterm birth in humans (44). Thus, elevated FM *COX2 in vivo* validated our *in vitro* data showing elevated FM PGE2 in response to methadone. There was also a trend for methadone exposure increasing mouse FM *IL1B* and *TNFA*, again, aligning with our *in vitro* findings.

Using non-pregnant rat and mouse *in vivo* models, rat, mouse and human *in vitro* cell models, and *in silico* approaches methadone has been shown to activate TLR4 by interacting with its adapter protein, MD2 (26). However, in our human FM explant model we did not find a role for TLR4/MD2 in mediating methadone-induced inflammation, which was in contrast to our findings for buprenorphine where TLR4/MD2 had a limited role (18). We instead found a dominant role for the μ-opioid receptor in mediating all inflammatory responses, with the exception of IL-1β that was NLRP3 inflammasome mediated, similarly to buprenorphine (18). Interestingly, while NLRP3 was only partially responsible for the methadone-induced FM IL-1β response, we did not find a role for another major inflammasome, AIM2. Thus, another inflammasome sensor may be contributing to methadone-induced IL-1β secretion. In alignment with our findings for a role for the μ-opioid receptor in mediating methadone-induced FM inflammation, the opioid receptor antagonist, (-) naloxone, similarly reduced all FM inflammatory responses to methadone, with the exception of IL-1β. The μ-opioid receptor is the primary target for opiate agonists (45) and can activate downstream inflammatory p38 MAPK and ERK signaling (33). Indeed we found that similarly to buprenorphine (18), methadone activates the p65 NFkB, p38 MAPK, ERK, and JNK signaling pathways, and these intermediate signaling pathways together mediate the methadone-induced FM inflammatory secretome.

Additional intermediate signals can contribute to the modulation and fine tuning of specific receptor-agonist mediated responses. We recently reported in human FMs that downstream of TLR2 activation by Gram-positive bacterial peptidoglycan, TLR4 activation by Gram-negative bacterial lipopolysaccharide, and TLR3 activation by viral dsRNA (Poly I:C), the TLR7/TLR8-activating miR-146a-3p (21, 28), acts as an endogenous DAMP that subsequently activates TLR7 and TLR8, contributing to the FM inflammasome-dependent and -independent inflammatory response (20, 29, 30). Herein, we identified this signaling pathway as playing a role in methadone-induced FM inflammation. While only FM TNF-α secretion in response to methadone was dependent on TLR7, all FM inflammatory factors elevated by methadone, with the exception of IL-1β, were TLR8-dependent. Since methadone increased human FM expression of miR-21a, this TLR7/TLR8-activating miR (31) may act as intermediate secondary signal contributing to opioid-driven FM inflammation.

While this mechanistic study included the use of primary human tissue explants and findings were complimented using a mouse model, we acknowledge that one limitation is the *in vitro* and *in vivo* systems modelled only acute opioid exposure. However, an important strength of our study is that we found most inflammatory responses triggered in human FMs by methadone could be mitigated by the opioid receptor antagonist, naloxone. Although naloxone is typically used as an emergency treatment for opioid overdose (27), it is safe to use in pregnancy, and studies indicate that buprenorphine-naloxone treatment during pregnancy may be more effective at lowering OUD relapse risk than buprenorphine monotherapy (46) and may reduce risk for NOWS and fetal growth restriction (47, 48). Combination buprenorphine-naloxone is the preferred treatment for OUD outside of pregnancy (48). Despite naloxone not being able to reduce methadone triggered IL-1β in human FMs, a known potent inducer of preterm birth, there still may be benefit in exploring combination methadone-naloxone therapy in pregnant women with OUD. Furthermore combining this dual therapy with an inflammasome inhibitor warrants investigation, and lab-based studies are underway.

In summary, methadone triggers an inflammatory response in both human and mouse FMs. In human FMs this inflammatory response is mediated by innate immune TLR7, TLR8 and NLRP3 signaling, the μ-opioid receptor, and downstream NFκB and ERK/JNK/MAPK signaling. Furthermore, the human FM inflammatory response to methadone can be mostly reduced by naloxone. Our findings advance our understanding of the mechanisms by which opioids may elevate the risk of preterm birth, and may help guide clinicians to optimize opiate agonist and antagonist treatments in pregnant women with OUD.

## ACKNOWLEDGMENTS

The authors would like to thank the Yale University Reproductive Sciences (YURS) Biobank and the staff of Labor and Delivery for tissue collection.

