## Supplemental Figure 1 for "Methadone Triggers Fetal Membrane Sterile Inflammation Through Innate Immune and Opioid Receptor Signaling"

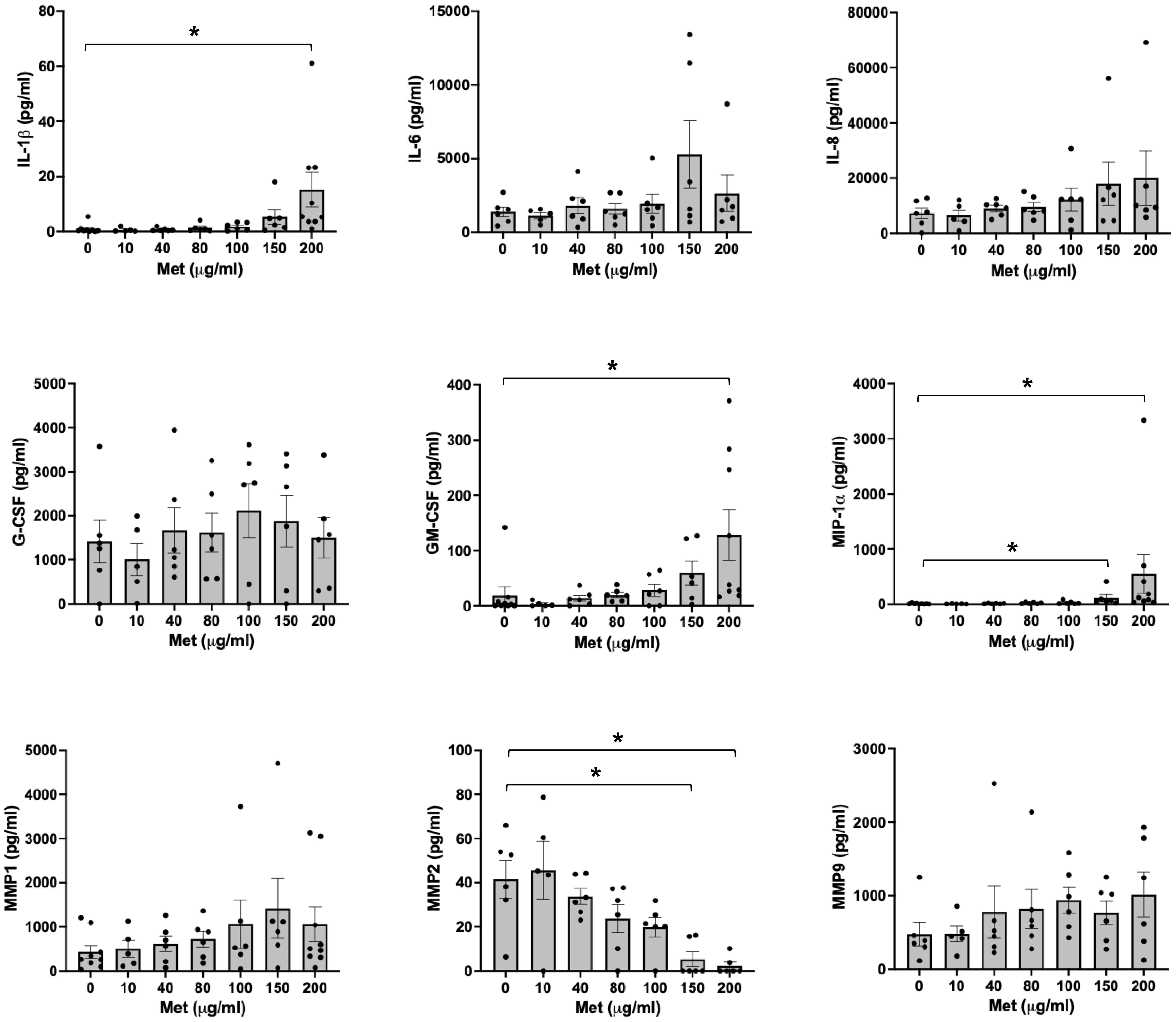


**Supplemental Figure 1. Methadone induces human FM sterile inflammation in a dose-dependent manner.** Human FM explants were treated with methadone (met) at 0, 10, 40, 80, 100, 150, or 200μg/ml (n=6-9). After 48hrs, cell-free supernatants were collected and measured by ELISA for IL-1β, IL-6, IL-8, TNFα, G-CSF, GM-CSF, MIP-1α, PGE2, MMP1, MMP2 and MMP9 (*$p$<0.05).
