## Supplemental Figure 2 for "Methadone Triggers Fetal Membrane Sterile Inflammation Through Innate Immune and Opioid Receptor Signaling"

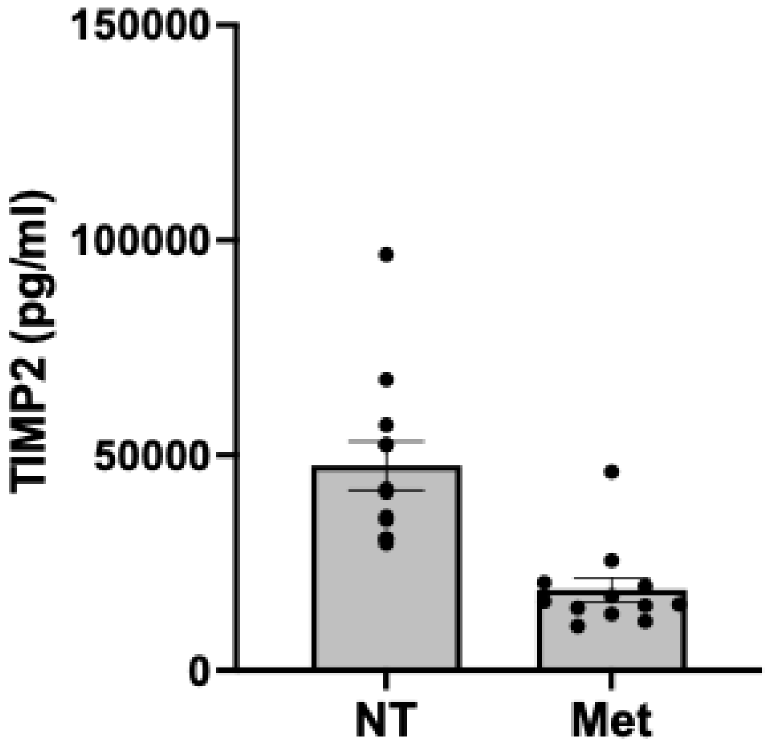


**Supplemental Figure 2. Methadone has no effect on human FM TIMP2 secretion.** Human FM explants were treated with no treatment (NT) or Methadone (Met) at 200μg/ml (n=12). After 48hrs, cell-free supernatants were collected and measured by ELISA for TIMP2.
