## Supplemental Figure 3 for "Methadone Triggers Fetal Membrane Sterile Inflammation Through Innate Immune and Opioid Receptor Signaling"

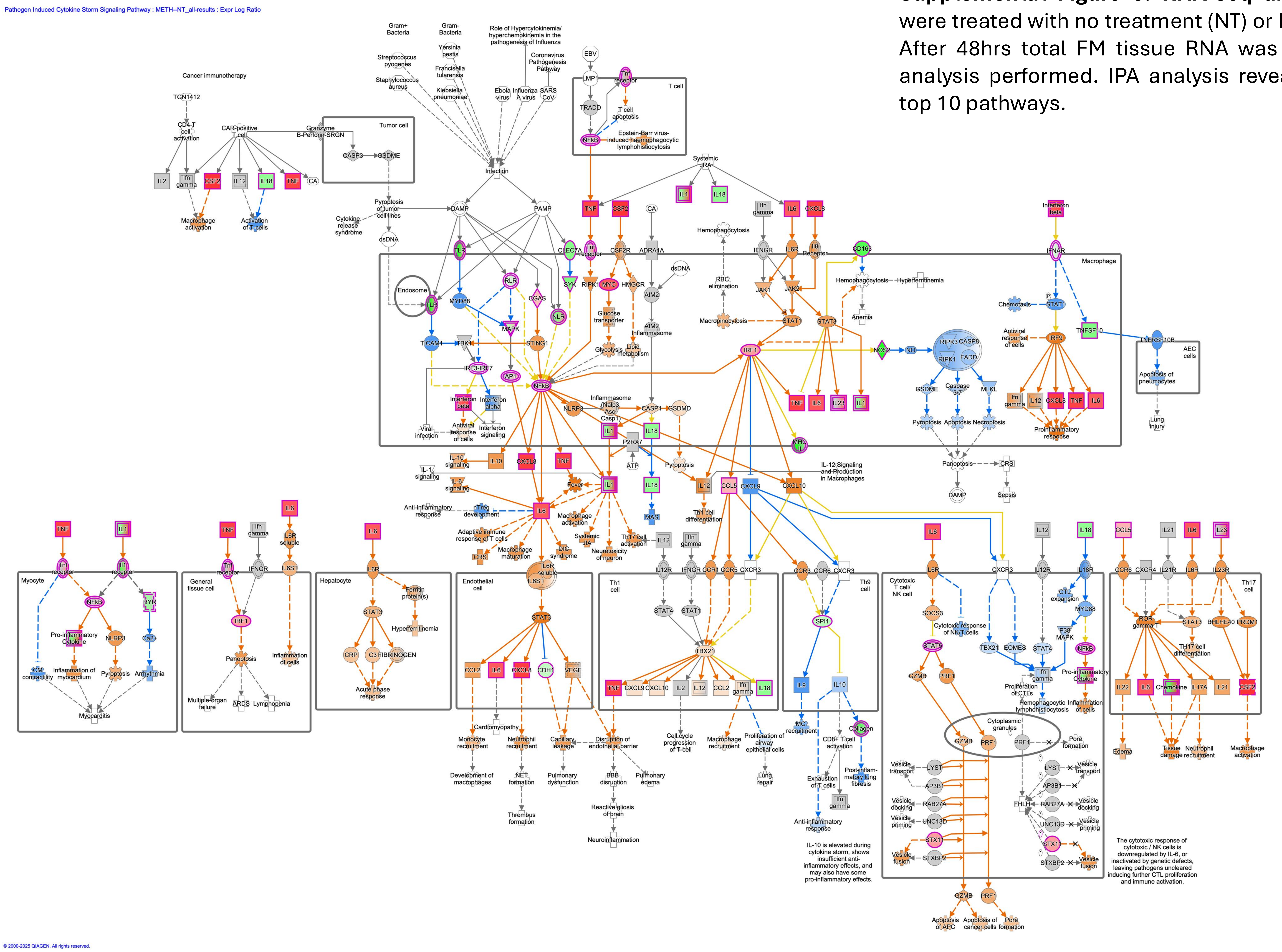


**Supplemental Figure 3. RNA-seq analysis.** Human FM explants were treated with no treatment (NT) or Methadone at 200µg/ml (n=6). After 48hrs total FM tissue RNA was extracted and bulk RNA-seq analysis performed. IPA analysis revealed “Cytokine Storm” in the top 10 pathways.
